# Branched chain amino acid aminotransferase 2 Regulates Ferroptotic Cell Death in Cancer Cells

**DOI:** 10.1101/2020.02.17.952754

**Authors:** Kang Wang, Zhengyang Zhang, Tsai Hsiang-i, Yanfang Liu, Ming Wang, Lian Song, Xiongfeng Cao, Zhanxue Xu, Hongbo Chen, Aihua Gong, Dongqing Wang, Fang Cheng, Haitao Zhu

**Author notes:** Correspondence to: Haitao Zhu, Fang Cheng, and Dongqing Wang. E-mail addresses (H. Zhu), (F. Cheng), (D. Wang). Kang Wang, Zhengyang Zhang and Tsai Hsiang-i contribute equally to this manuscript.

## Abstract

Ferroptosis has been implicated as a tumor-suppressor function for cancer therapy. Recently the sensitivity to ferroptosis was tightly linked to numerous biological processes, including metabolism of amino acid. Here, using a high-throughput CRISPR/Cas9 based genetic screen in HepG2 cells to search for metabolic proteins inhibiting ferroptosis, we identified branched chain amino acid aminotransferase 2 (BCAT2) as a novel suppressor of ferroptosis. Mechanistically, ferroptosis inducers (erastin, sorafenib and sulfasalazine) activated AMPK/SREBP1 signaling pathway through ferritinophagy, which in turn inhibited BCAT2 transcription. We further confirmed that BCAT2 mediating the metabolism of sulfur amino acid, regulated intracellular glutamate level, whose activation by ectopic expression specifically antagonize system Xc– inhibition and protected liver and pancreatic cancer cells from ferroptosis in vitro and in vivo. Finally, our results demonstrate the synergistic effect of sorafenib and sulfasalazine in downregulating BCAT2 expression and dictating ferroptotic death, where BCAT2 can also be used to predict the responsiveness of cancer cells to ferroptosis-inducing therapies. Collectively, these findings identify a novel role of BCAT2 in ferroptosis, suggesting a potential therapeutic strategy for overcoming sorafenib resistance.

## Introduction

Ferroptosis is emerging as an iron-dependent type of regulated cell death, which is induced by the loss of cellular redox homeostasis, leading to unchecked lipid peroxidation and eventually cell death(Dixon, Lemberg et al., 2012). Pharmacological inactivation of cystine-glutamate antiporter (system Xc^−^), or glutathione peroxidase 4 (GPX4) can induce ferroptosis(Dixon, 2017), suggesting the crucial roles of the glutathione-dependent antioxidant defenses in preventing ferroptotic cell death.

Ferroptosis has been implicated in ischemia-induced organ injury, pathological cell death associated with degenerative diseases, and in different types of cancer (Friedmann Angeli, Schneider et al., 2014, Hambright, Fonseca et al., 2017, Lu, Chen et al., 2017). A variety of tumor cells are susceptible to ferroptosis including lymphoma, cervical cancer, head and neck cancer, pancreatic cancer, renal cell carcinoma and hepatocellular carcinoma (HCC). Various studies have confirmed the pivotal role of ferroptosis inducers, including small-molecule ferroptosis inducers such as erastin as well as a number of drugs (eg. sorafenib, artemisinin and its derivatives) in killing tumor cells and suppressing tumor growth(Dixon, Patel et al., 2014, Ooko, Saeed et al., 2015, Xie, Hou et al., 2016). These ferroptosis inducers also synergy with chemotherapeutic drugs in cancer treatment. Interestingly, some types of cancer are more sensitive to ferroptosis inducers than others. The reverse transsulfuration pathway has been identified as a negative regulator of ferroptosis and a deficiency of this pathway in ovarian cancer cells are associated with increased sensitivity to erastin-induced ferroptosis(Liu, Lin et al., 2019). HSF1-HSPB1 pathway also negatively regulated erastin-induced ferroptosis in human cervical cancer, prostate cancer and osteosarcoma(Sun, Ou et al., 2015). MUC1-C/xCT pathway is another negative regulator in erastin-induced ferroptosis of triple-negative breast cancer cells(Hasegawa, Takahashi et al., 2016).

Accumulating evidence indicates that cellular metabolism plays a crucial role in ferroptosis (Angeli, Shah et al., 2017, Stockwell, Friedmann Angeli et al., 2017). The transcriptional factor NRF2 coordinates the antioxidant defensive system in the regulation of ferroptosis. The p62-Keap1-NRF2 is a central inhibitory pathway of ferroptosis in liver cancer cells (Sun, Ou et al., 2016). Genetic or pharmacologically inhibition of NRF2 significantly enhanced ferroptosis susceptibility of liver cancer induced by erastin and sorafenib, whereas the activation of NRF2 expression led to cellular resistance to ferroptosis, suggesting a central role of the partially reduced oxygen-containing molecules, especially reactive oxygen species (ROS) in ferroptosis. Intracellular iron metabolism is also essential for ferroptosis through either the action of iron-dependent oxidases, or by Fenton chemistry. A recent report suggests autophagic degradation of ferritin regulates ferroptosis through an autophagy cargo receptor NCOA4 (Hou, Xie et al., 2016). Not surprisingly, amino acid metabolism is also involved in ferroptosis (Gao, Monian et al., 2015, Hayano, Yang et al., 2016).

High concentration of extracellular glutamate, erastin, or other system Xc^−^ inhibitors block intracellular cystine/cysteine uptake to induce ferroptosis. Silencing cysteinyl-tRNA synthetase (CARS) upregulates the transsulfuration pathway, which lead to resistance to erastin-induced ferroptosis. Glutamine mediates ferroptosis through its specific metabolic enzymes, glutaminases (GLS1 and GLS2), though the mechanism of glutaminolysis process is complex. However, the metabolic pathways controlling ferroptosis sensitivity in liver cancer cells remains unclear.

In this study, we identify a branched-chain amino acid aminotransferase 2 (BCAT2), an aminotransferase enzyme mediating sulfur amino acid metabolism, as a specific inhibitor of ferroptosis. We show that BCAT2 is involved in system Xc^−^ inhibitor induced ferroptosis in liver cancer cells. Furthermore, BCAT2 participates in the mechanisms for sulfasalazine synergizing with sorafenib to induce ferroptosis. Thus, our results demonstrate that BCAT2 serves as a suppressor of ferroptosis, and contributes to the core metabolic signaling pathways involved in liver cancer ferroptosis.

## Results

### Identification of novel players of ferroptosis by kinome CRISPRa screening

Ferroptosis can be induced by two classes of small-molecule substances known as class 1 system Xc^−^ inhibitors (including erastin, sulfasalazine, and sorafenib) and class 2 ferroptosis inducers (GPX4 inhibitors). We first tested and confirmed the effects of erastin (Erastin), sorafenib (SOR) and sulfasalazine (SAS) as probes to induce ferroptosis in human pancreatic cancer cell line Aspc-1, human hepatocellular carcinoma cell line HepG2, human colorectal cancer cell line SW480, as well as human fibrosarcoma cell line HT1080. The results confirmed that erastin, sorafenib and sulfasalazine could significantly induce the cancer cell death at the concentration at 10 μmol/L for erastin, 5 μmol/L for sorafenib and 1 mmol/L for sulfasalazine, respectively (Figure S1A and S1B). Furthermore, the cell death was dramatically inhibited by ferroptosis inhibitor ferrostin-1, but not by apoptotic inhibitor ZAVD-FMK or necroptosis inhibitor Necrosulfonamide, indicating the specificity of all three ferroptosis inducers ((Figure S1A and S1B).

To systemically elucidate conserved downstream negative regulators of ferroptosis, we performed a large-scale genetic CRISPR activation (CRISPRa) screen. A pooled human CRISPRa sgRNA lentivirus library targeting 2320 Kinases, Phosphatases, and Drug Targets (https://www.addgene.org/pooled-library/weissman-human-crispra-v2-subpools/) together with Cas9-VPR enzyme were introduced into HepG2 cells by lentiviral transduction, which were then treated with erastin or control DMSO (Figure 1A). Deep sequencing of the sgRNAs integrated into genomic DNA from control cells and cells that survived ferroptosis induction was subsequently performed. Comparison of the sequencing data led to the identification of sgRNAs that were enriched in cells surviving ferroptosis treatment. The gene targets of the enriched sgRNAs are potential genes that confer resistance to erastin-mediated ferroptosis in HepG2 cells. Among the screen hits, many reported ferroptosis genes were identified and validated in our screen approach. Intriguingly, our screen also identified genes not previously implicated in the regulation of ferroptosis, among which BCAT2 was the top candidate of potential negative regulators in ferroptosis (Figure 1A).

**Figure 1.**
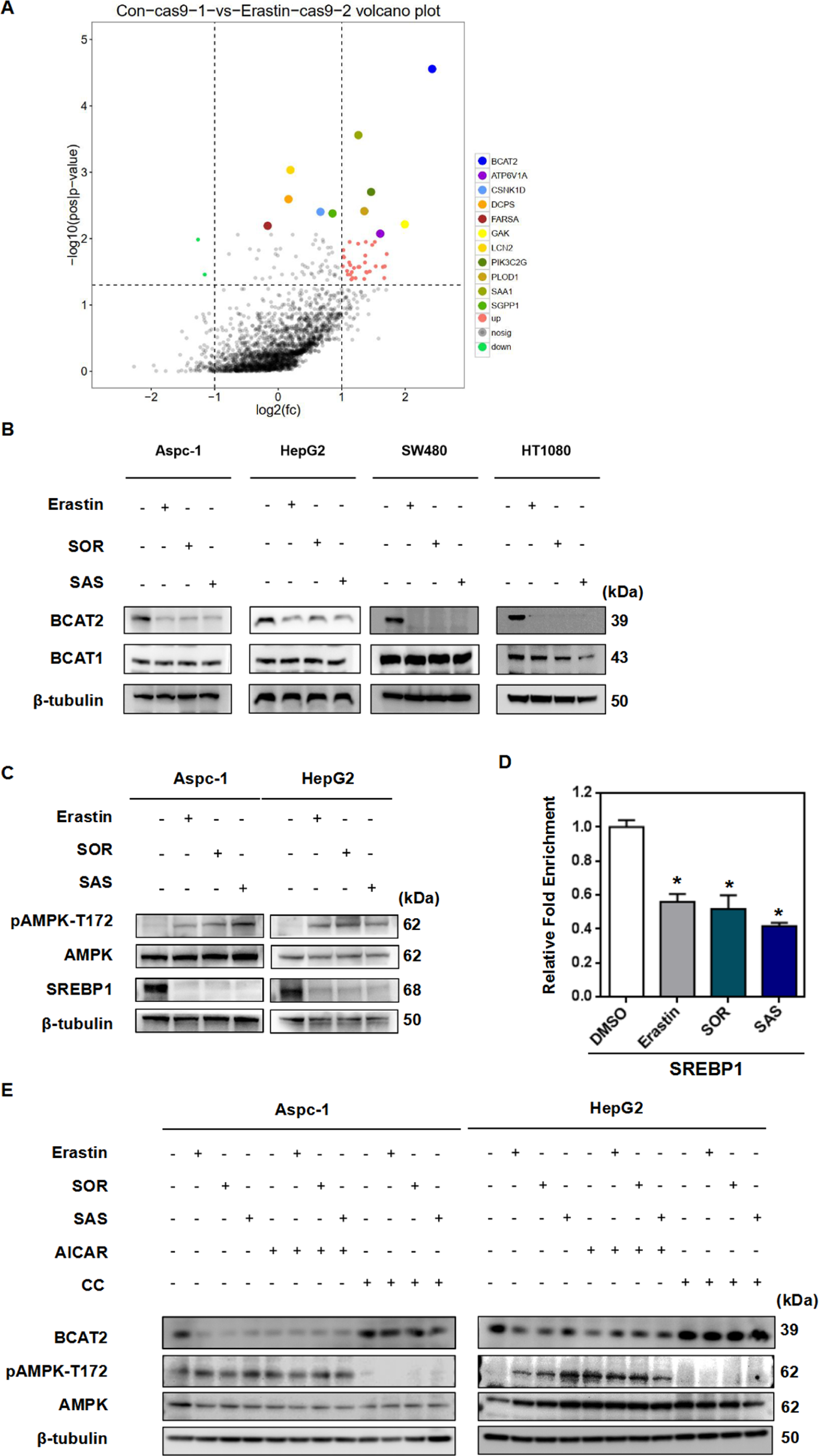
Ferroptosis inducers inhibit BCAT2 expression through ferrtinophagy-AMPK-SREBP1 pathway. A A large-scale genetic CRISPR activation (CRISPRa) screen identifies genes essential for regulating ferroptosis. HepG2 cells expressing dcas9 were mutagenized with a pooled lentiviral sgRNA library. Significant hits from screens in cells treated with erastin or DMSO-treated cells. Dots represent individual genes. Colorful dots indicate significant enrichment genes that are resistant to erastin mediated ferroptosis. X axis indicated the fold change of gRNA insertions per gene (treatment group over control group), Y axis represents the frequency of insertions (*p*<0.05). B Western blot analysis the protein expression levels of BCAT2 and BCAT1 in Aspc-1, HepG2, SW480 and HT1080 cells treated with erastin (20 or 10 μmol/L), sorafenib(10 or 5 μmol/L) or sulfasalazine(2 or 1 mmol/L). β-tubulin expression was detected as a loading control. C Western blot analysis the protein expression levels of AMPK, pAMPK(T172), and SREBP1 in Aspc-1 and HepG2 cells treated with DMSO (control), erastin (20 or 10 μmol/L), sorafenib(10 or 5 μmol/L) or sulfasalazine(2 or 1 mmol/L). β-tubulin expression was detected as a loading control. D Chromatin immunoprecipitation (ChIP) analysis of SREBP1 binding to BCAT2 promoter in HepG2 cells treated with DMSO (control), erastin (20 μmol/L), sorafenib(10 μmol/L) or sulfasalazine(2 mmol/L). E Western blot analysis the protein expression levels of BCAT2, AMPK, and pAMPK(T172) in Aspc-1, and HepG2 cells treated with DMSO (control), erastin (20 μmol/L), sorafenib(10 μmol/L) or sulfasalazine(2 mmol/L) in the absence or presence of AICAR (AMPK activator, 2 mmol/L) and Compound C (AMPK inhibitor, 1 μmol/L). β-tubulin expression was detected as a loading control. Data information: Experiments were repeated three times, and the data are expressed as the mean ± SEM. \**p* < 0.05, vs. control group. Statistical analysis was performed using Student’s t-test.

Branched-chain amino acid aminotransferase (BCAT) is an aminotransferase enzyme which acts upon branched-chain amino acids (BCAAs) to regulate sulfur amino acid metabolism. To validate the role of BCAT2 in ferroptosis, we first investigated the expression of BCAT2 in above mentioned four cancer cell lines upon induction of erastin, sorafenib, and sulfasalazine. Western blot and qRT-PCR showed that in all four cell lines, there was a reduction of BCAT2, but not BCAT1 (a paralog of BCAT2) in protein- and mRNA-expression levels upon ferroptosis inducer treatment, which was reversed in the presence of deferoxamine mesylate(DFO, a ferroptosis inhibitor) (Figure 1B, Figure S2A, S2B and S6). BCAT2 reduction was reversed in the presence of DFO (Figure 1B, Figure S2A, S2B and S6). Moreover, erastin, sorafenib, and sulfasalazine downregulated the BCAT2 protein level in a time dependent manner (Figure S2C). TCGA data analysis showed that BCAT2 expression correlated with cancer grade (Figure S3A) and the expression of ferroptosis markers (GPX4 and NCOA4) in hepatocellular carcinoma (Figure S3B).

### Ferroptosis inducers inhibit BCAT2 expression through ferritinophagy-AMPK-SREBP1 pathway

It has been reported that AMP-activated protein kinase (AMPK) inhibits nuclear translocation of sterol response element binding protein 1 (SREBP1), which consequently suppresses the transcription of its direct target gene BCAT2 (Dey, Baddour et al., 2017). Therefore, we hypothesize that ferroptosis inducers downregulate BCAT2 via AMPK-SREBP1 signaling pathway. We confirmed that erastin, sorafenib or sulfasalazine induced AMPK phosphorylation on threonine residue 172 (T172) and reduced the expression of SREBP1, assessed by quantifying the signals from western blotting (Figure 1C, Figure S4A and S4B). ChIP assay also revealed that SREBP1 binding to BCAT2 was significantly reduced in the presence of erastin, sorafenib, or sulfasalazine (Figure 1D), suggesting they further prevent the transcription factor SREBP1 to activate BCAT2 transcription in the nucleus. Moreover, AMPK activator AICAR downregulated BCAT2 expression in both Aspc-1 and HepG2 cancer cells in a similar manner to erastin, sorafenib or sulfasalazine, which can be reversed by AMPK inhibitor Compound C, further confirming that ferroptosis inhibitors downregulate BCAT2 expression through activating AMPK (Figure 1E, Figure S4C and S4D). Collectively, these results indicate that AMPK/SREBP1 mediates BCAT2 expression in ferroptotic process.

Next, we would like to understand how AMPK/SREBP1 is activated by ferroptosis inducers. As AMPK is a promoter of autophagy and ferroptosis is an autophagic cell death process called ferritinophagy (Gao, Monian et al., 2016, Song, Zhu et al., 2018), we asked whether ferritinophagy is involved in AMPK activity. Nuclear receptor coactivator 4 (NCOA4) was a selective cargo receptor for the selective autophagic turnover of ferritin (Hou et al., 2016). First, we found that erastin, sorafenib or sulfasalazine inhibited NCOA4 expression (Figure S5A) and the formation of GFP-LC3 puncta, the hallmarks of autophagy response (Figure S5B), whereas inhibition of NCOA4 or autophagy can further increase Fe^2+^ level (Figure S5C-F). Moreover, erastin-induced the AMPK phosphorylation on threonine residue 172 (T172) and BCAT2 inhibition can be reversed in the presence of BafA1 (an autophagy inhibitor) and DFO (an iron chelator) (Figure S6), further indicating AMPK/SREBP1 is the downstream of ferritinophagy and ferrous ions to inhibit BCAT2 expression. Therefore, erastin, sorafenib or sulfasalazine inhibits BCAT2 expression through ferritinophagy -AMPK-SREBP1 pathway.

### BCAT2 is a suppressor of ferroptotic cancer cell death

To investigate the role of BCAT2 in ferroptosis, we first transfected BCAT2 cDNA plasmid into Aspc-1 and HepG2 cells and confirmed the overexpression of BCAT2 proteins in these cells by western blot (Figure S7A). Given the critical role of iron in ferroptosis, we first examined the correlation of BCAT2 expression and iron accumulation. As expected, erastin, sorafenib, or sulfasalazine treatment induced free iron accumulation in both control and BCAT2 transfected cells (Figure 2A and Figure S7B). Compared to the parental cells, overexpression of BCAT2 had no effect on the accumulation of Fe^2+^ in Aspc-1 and HepG2 cells following ferroptotic induction (Figure 2A and Figure S7B). Because system Xc^−^ is responsible for maintaining redox homeostasis by importing cysteine to synthesize GSH, we asked whether BCAT2 is involved in system Xc^−^ mediated GSH activity. GSH level was inhibited in Aspc-1 and HepG2 cells following erastin, sorafenib, and sulfasalazine treatment, which was restored by ectopic expression of BCAT2 (Figure 2B and Figure S7C). The level of malondialdehyde (MDA), an end product of lipid peroxidation, was increased in Aspc-1 and HepG2 cells following erastin, sorafenib, and sulfasalazine treatment, but decreased in BCAT2 overexpressing cells compared to their parental cells (Figure 2C and Figure S7D). In line with these results, in the presence of erastin, sorafenib, or sulfasalazine, BCAT2 overexpression increased intracellular glutamate ((Figure 2D and Figure S7E) and the glutamate release ((Figure 2E and Figure S7F), and reduced system Xc^−^ inhibitor-induced cell death (Figure 2F and Figure S7G) in a time-dependent manner. In order to confirm the relationship between BCAT2 and ferroptosis *in vivo*, a subcutaneous xenograft tumor model was established by injecting 1 × 10^6^ parental or overexpression BCAT2 Panc02 cancer cells into the nude mouse. Administration of erastin into the mice reduced the size of Panc02 parental tumors by 64%, compared with vehicle-treated tumors at day 14 (Figure S7H). In agreement with *in vitro* results, BCAT2 protected Panc02 cancer cells against erastin-induced reduction in tumor growth by 2.5-fold (Figure S7I), indicating that overexpression BCAT2 rescues erastin induced tumor inhibition.

**Figure 2.**
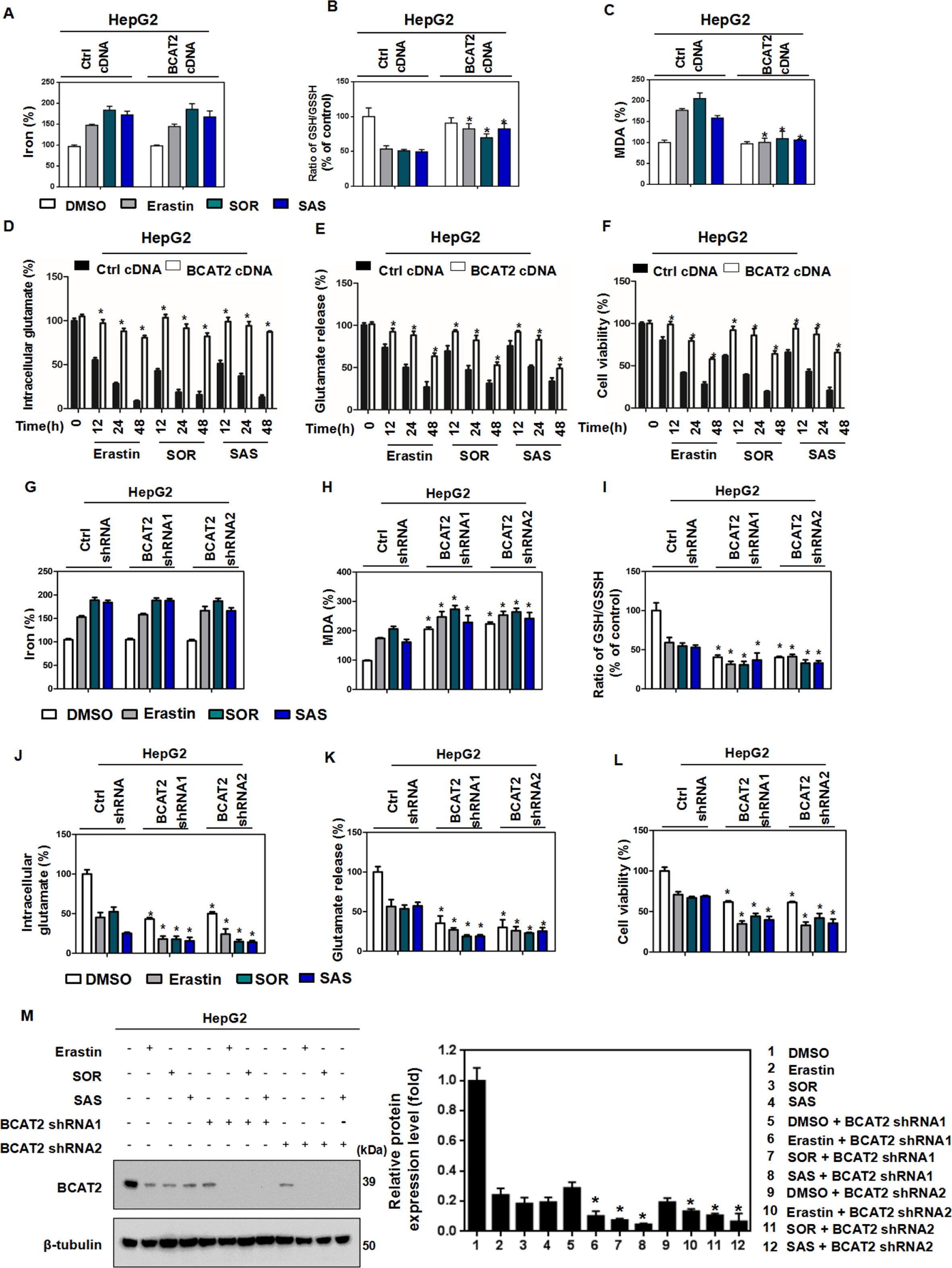
BCAT2 is a negative regulator of ferroptotic cancer cell death. A-C BCAT2 overexpressed and parental HepG2 cells were treated with DMSO (control), erastin (20 μmol/L), sorafenib(10 μmol/L) or sulfasalazine(2 mmol/L). The relative levels of Fe^2+^ (A), ratio of GSH/GSSH (B), and MDA (C) were assayed. D-F BCAT2 overexpressed and parental HepG2 cells were treated with DMSO (control), erastin (20 μmol/L), sorafenib(10 μmol/L) or sulfasalazine(2 mmol/L) for the indicated time(0, 12, 24, 48 h). The relative levels of intracellular glutamate(D), glutamate release(E) and cell viability(F) were assayed. G-L BCAT2 knockdown and parental HepG2 cells were treated with DMSO (control), erastin (20 μmol/L), sorafenib(10 μmol/L) or sulfasalazine(2 mmol/L). The relative levels of Fe^2+^ (G), MDA (H), ratio of GSH/GSSH (I), intracellular glutamate (J), glutamate release (K) and cell viability (L) was assayed. M Western blot analysis of the protein expression levels of BCAT2 in BCAT2-knockdown and parental HepG2 cells treated with or without Erastin (20 μmol/L), sorafenib(10 μmol/L) or sulfasalazine(2 mmol/L). β-tubulin expression was detected as a loading control. Data information: Experiments were repeated three times, and the data are expressed as the mean ± SEM. \**p* < 0.05 vs. control group. Statistical analysis was performed using Student’s t-test.

To further elucidate the functional role of BCAT2 in ferroptosis, two stable BCAT2-knockdown cell clones (BCAT2 shRNA1 and shRNA2) were established with high silencing efficiency (up to 60% silencing) confirmed by western blotting (Figure S8A). Compared to the parental cancer cells, BCAT2 knockdown cells showed smaller mitochondria morphology with more condensed mitochondrial membrane density, and reduced mitochondria crista, which are typical morphological features of ferroptosis (Figure S8B). Furthermore, knockdown of BCAT2 significantly increased MDA production (Figure 2H and Figure S8D) and GSH depletion (Figure 2I and Figure S8E) in Aspc-1 and HepG2 cells in the presence of erastin, sorafenib, and sulfasalazine, but had no effect on accumulation of free cellular iron (Figure 2G and Figure S8C). Moreover, BCAT2 knockdown decreased the level of intracellular glutamate (Figure 2J and Figure S8F), the glutamate release (Figure 2K and Figure S8G) as well as cell viability (Figure 2L and Figure S8H) in the presence of erastin, sorafenib, or sulfasalazine. Accordingly, the colony formation capability got inhibited in BCAT2 silencing cells (Figure S8I). Moreover, knockdown of BCAT2 did not affect the SLC7A11 and GPX4 protein expression level in Aspc-1 and HepG2 cells (Figure S9A). These results demonstrated that BCAT2 knockdown could partly induce the ferroptosis.

### BCAT2 participates in the mechanisms for sulfasalazine synergizing with sorafenib to induce ferroptosis

As sorafenib or sulfasalazine suppressed BCAT2 knockdown in a similar manner, we hypothesized that sorafenib or sulfasalazine may have synergistic effect in inducing ferroptosis. We first investigated the role of combining sorafenib and sulfasalazine in BCAT2 expression. Intriguingly, we found that combination sorafenib and sulfasalazine dramatically inhibited BCAT2 expression in Aspc-1 and HepG2 cells (Figure 3A), in a comparable pattern to sorafenib or sulfasalazine alone, together with BCAT2 shRNA (Figure 2M and S8J). Sorafenib and sulfasalazine also exhibited a synergistic effect in increasing the cell death (Figure 3B) and MDA production (Figure 3C), suppressing the glutamate release (Figure 3D) and the intracellular glutamate level (Figure 3E), which can be rescued in the presence of ferrostatin-1. All these results support our hypothesis that the effect of sorafenib and sulfasalazine on ferroptosis partially through regulating BCAT2 expression.

**Figure 3.**
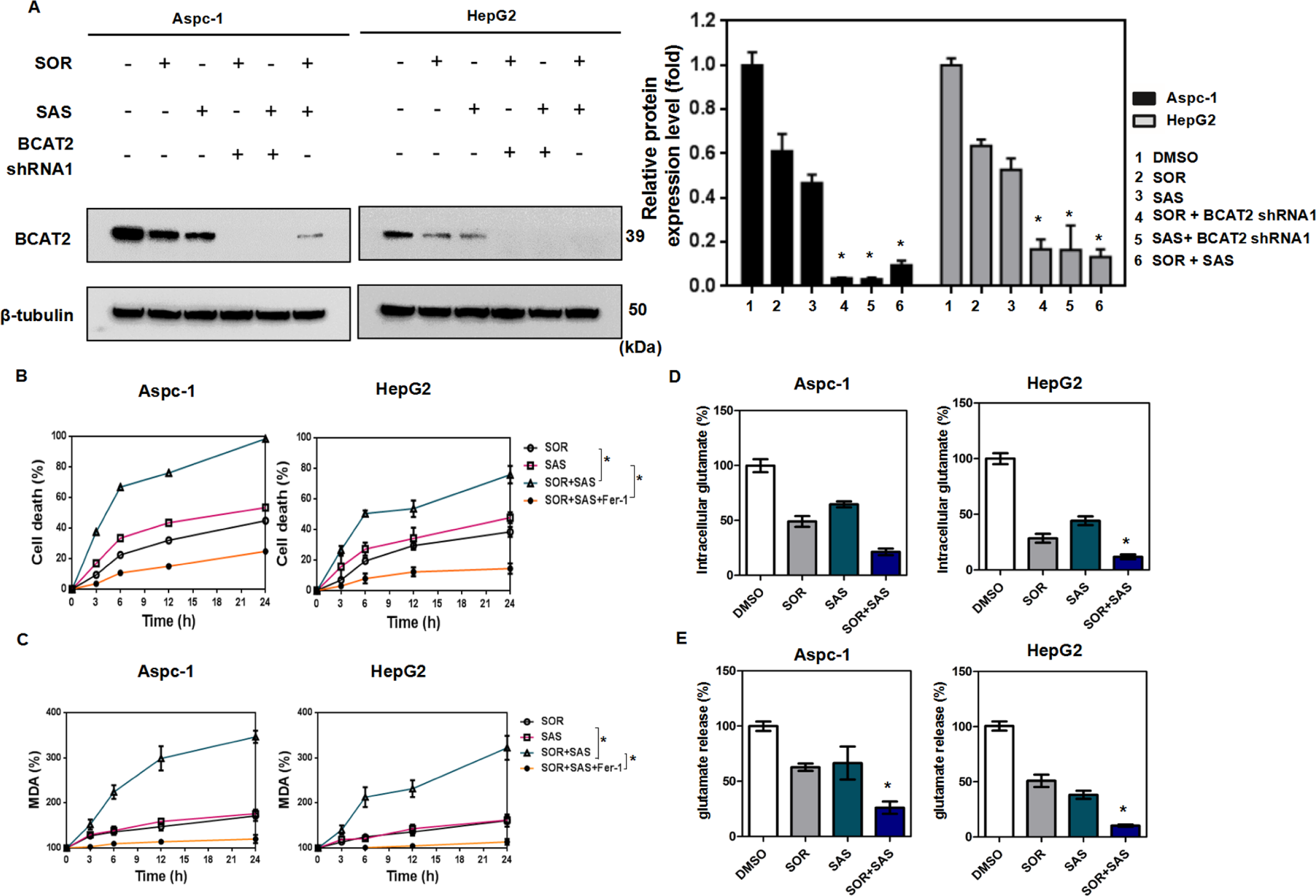
BCAT2 participates in the mechanisms for sulfasalazine synergizing with sorafenib to induce ferroptosis *in vitro*. A Western blot analysis of the protein expression levels of BCAT2 in BCAT2-knockdown and parental Aspc-1 and HepG2 cells treated with or without sorafenib(10 or 5 μmol/L), sulfasalazine(2 or 1 mmol/L) or sorafenib(10 or 5 μmol/L) + sulfasalazine(2 or 1 mmol/L). β-tubulin expression was detected as a loading control. B Aspc-1 and HepG2 cells were treated with sorafenib(10 or 5 μmol/L), sulfasalazine(2 or 1 mmol/L) or sorafenib(10 or 5 μmol/L) + sulfasalazine(2 or 1 mmol/L) in the absence or presence of ferrostatin-1(1 μmol/L) for the indicated time, and cell death was analyzed by propidium iodide staining. C Aspc-1 and HepG2 cells were treated with sorafenib(10 or 5 μmol/L), sulfasalazine(2 or 1 mmol/L) or sorafenib(10 or 5 μmol/L) + sulfasalazine(2 or 1 mmol/L) in the absence or presence of ferrostatin-1(1 μmol/L). The relative levels of MDA were assayed. D, E Aspc-1 and HepG2 cells were treated with DMSO (control), sorafenib(10 or 5 μmol/L), sulfasalazine(2 or 1 mmol/L) or sorafenib(10 or 5 μmol/L) + sulfasalazine(2 or 1 mmol/L). The relative levels of intracellular glutamate(D) and glutamate release (E) were assayed. Data information: Experiments were repeated three times, and the data are expressed as the mean ± SEM. \**p* < 0.05 vs. control group. Statistical analysis was performed using Student’s t-test.

We next investigated whether sorafenib and sulfasalazine has synergistic anticancer effect *in vivo*. Administration of sorafenib and sulfasalazine reduced the size of Panc02 subcutaneous tumors in C57BL/6 mice by 9.63% and 13.5%, respectively, and the combination therapy further reduced the size by 81.39%, compared with vehicle-treated tumors at day 14 (Figure 4A, 4B and 4C). Since orthotopic xenograft models are considered superior to the subcutaneous tumor models in terms of replicating the tumor microenvironment and predicting of drug efficacy, we would like to check whether induction of ferroptosis by sulfasalazine also enhances the anticancer activity of sorafenib in orthotopic hepatocellular carcinoma models with established mouse H22 cells in C57BL/6 mice(Figure 4D). Indeed, sorafenib combined with sulfasalazine significantly reduced the tumor size (Figure 4E) and prolonged animal survival (Figure 4F) in orthotopic xenograft tumor.

**Figure 4.**
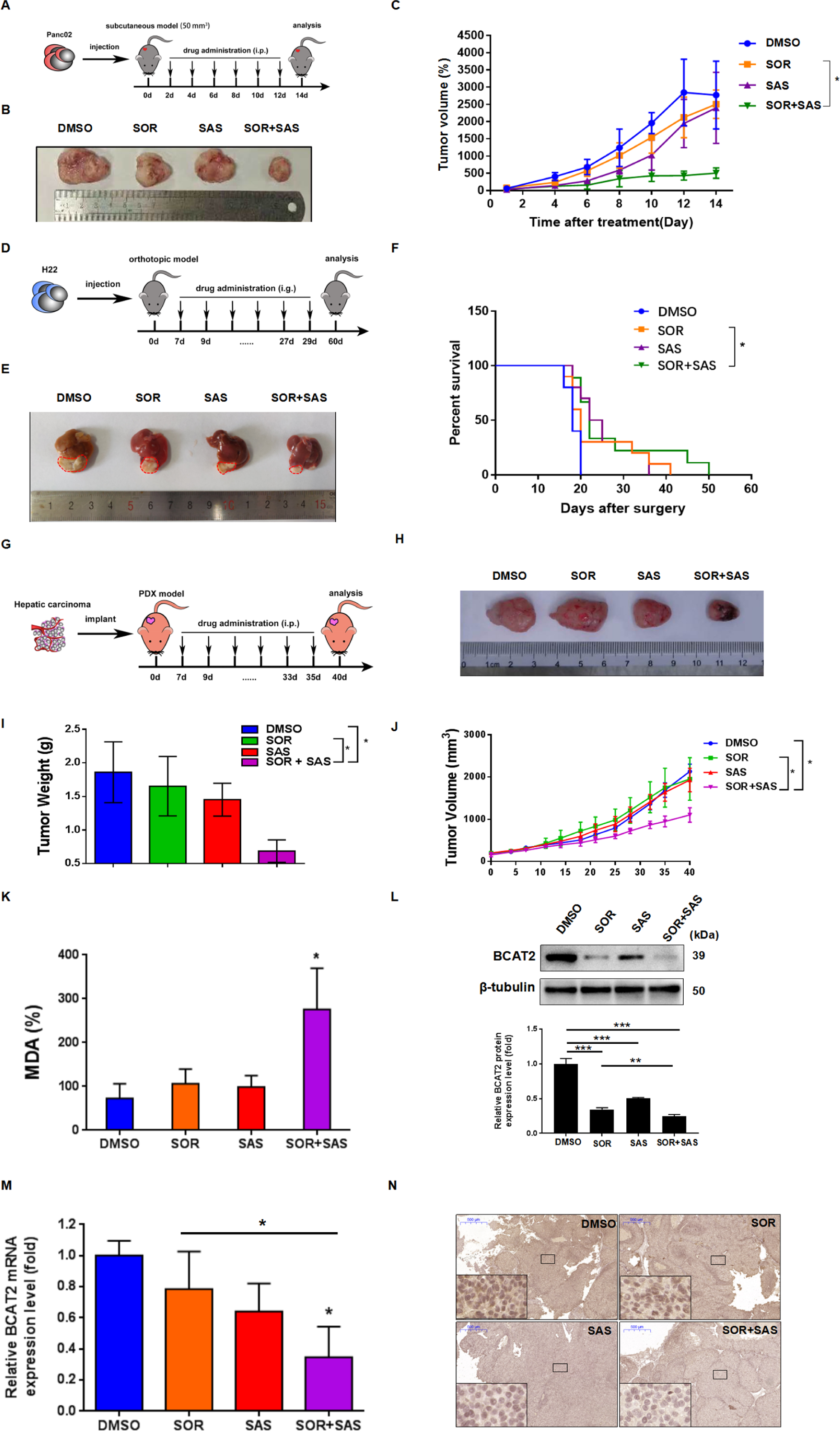
BCAT2 participates in the mechanisms for sulfasalazine synergizing with sorafenib to induce ferroptosis *in vivo*. A Schematic representation of the *in vivo* sorafenib and sulfasalazine combination anticancer effect in Panc02 subcutaneous tumor model. C57BL/6 mice were injected subcutaneously with 1 × 10^6^ Panc02 cancer cells and treated with DMSO(control group), sorafenib(10 mg/kg/i.p., every two days), sulfasalazine (100 mg/kg/i.p., every two days), or sorafenib(10 mg/kg/i.p., every two days) + sulfasalazine (100 mg/kg/i.p., every two days) for 2 weeks(n = 5 mice/group). B Representative photographs of isolated tumor tissues at day 14 after treatment. C Tumor volume was calculated every two days. D Schematic representation of the *in vivo* sorafenib and sulfasalazine combination anticancer effect in H22 orthotopic xenograft tumor model. 1 × 10^6^ H22 cells were injected into left lobe of C57BL/6 mice livers and following treated with DMSO(control group), sorafenib(30 mg/kg/i.g., every two days), sulfasalazine (100 mg/kg/i.g., every two days), or sorafenib(10 mg/kg/i.g., every two days) + sulfasalazine (100 mg/kg/i.g., every two days) for 2 weeks(n = 10 mice/group). E Representative photographs of isolated tumor tissues following various treatments. F Animal survival was calculated every day for 2 months (KaplanMeier survival analysis). G Schematic representation of established PDX hepatocellular carcinoma models were treated with DMSO(control group), sorafenib(10 mg/kg/i.p., every two days), sulfasalazine (100 mg/kg/i.p., every two days), or sorafenib(10 mg/kg/i.p., every two days) + sulfasalazine (100 mg/kg/i.p., every two days) for 2 weeks. H Representative photographs of isolated tumor tissues at day 40 after treatment. I Tumor volume was calculated every three days. J Tumor weight of isolated tumor tissues at day 40 after treatment. K MDA levels in isolated tumors at day 40 after treatment were assayed. L Western blot analysis of the protein expression level of BCAT2 in isolated tumor tissues at day 40 after treatment. M qRT-PCR analysis of mRNA expression level of BCAT2 in isolated tumor tissues at day 40 after treatment. N Immunohistochemistry analysis of the expression of BCAT2 in isolated tumor tissues at day 40 after treatment. Data information: Experiments were repeated three times, and the data are expressed as the mean ± SEM. * *p* < 0.05, ** *p* < 0.01, *** *p* < 0.001 vs. control group. Statistical analysis was performed using Student’s t-test.

**Figure 5.**
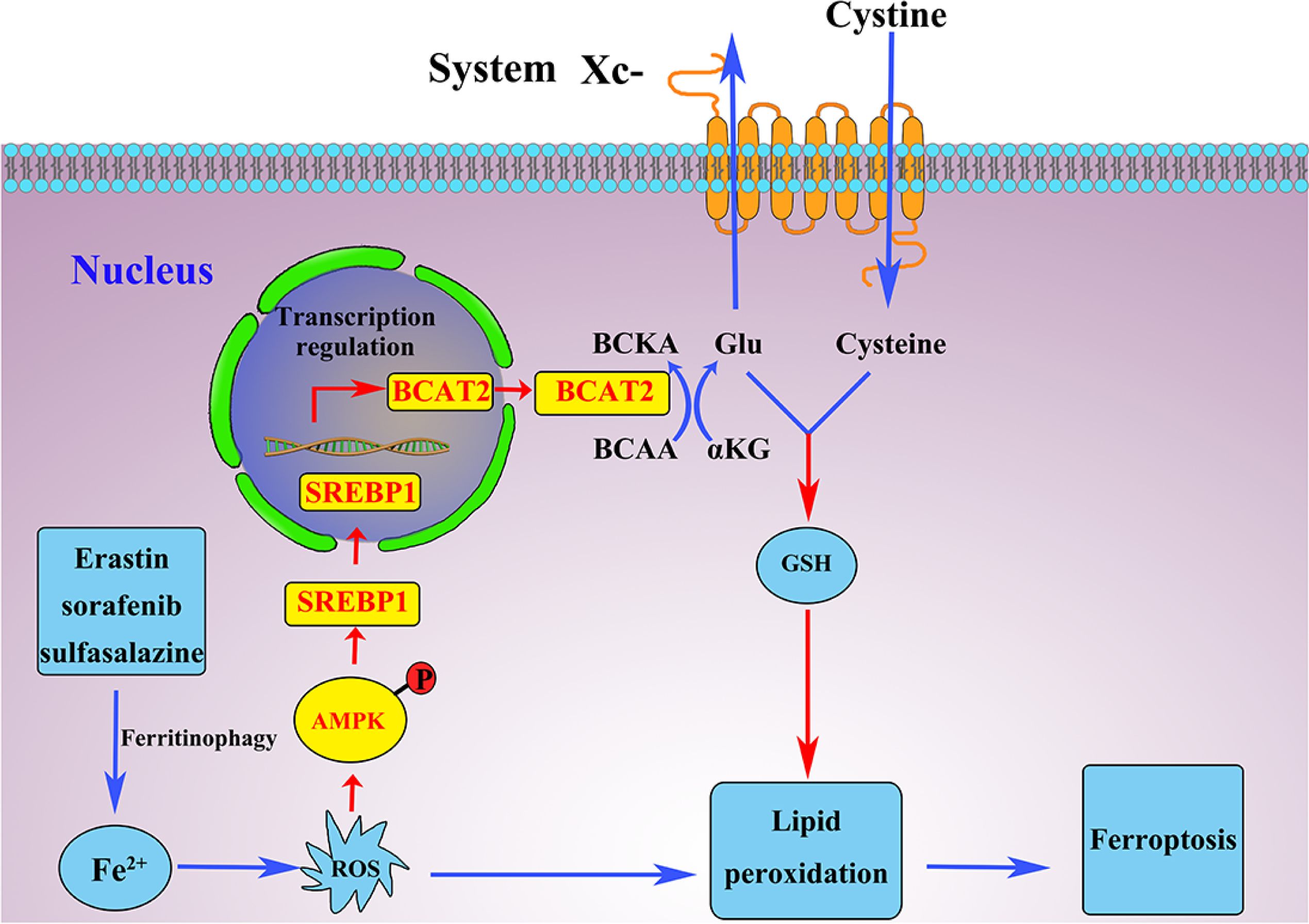
Schematic representation of the mechanisms of BCAT2 participating in sulfasalazine synergizing with sorafenib to induce ferroptosis.

To order to reveal more clinical relevance, we turned to a patient-derived xenografts (PDXs) model, which has been applied to pre-clinical drug testing in many types of cancers due to its biologically stable and accurately reflect the patient tumor with regards to histopathology, gene expression, genetic mutations, and therapeutic response (Figure 4G). In line with the previous results, sorafenib combined with sulfasalazine significantly reduced the tumor size by 48% and tumor weight by 63% in PDX model tumor (Figure 4H-J). Compared to the control group, sorafenib combined with sulfasalazine significantly augmented locally MDA levels (Figure 4K). Moreover, there was a significant reduction of BCAT2 in protein- and mRNA-expression upon combination treatment of sorafenib and sulfasalazine (Figure 4L and 4M). These *in vivo* results further support the in vitro evidence that collective inhibition of the BCAT2 pathway effectively enhances the anticancer activity by induction of ferroptosis.

## Discussions

In this study, we confirmed that erastin, sorafenib or sulfasalazine activate ferritinophagy and increase cellular labile iron level. High levels of cellular labile iron consequently lead to rapid accumulation of cellular ROS, which is essential for ferroptosis. Interestingly, we found this ferritinophagy pathway also activate AMPK phosphorylation, which consequently suppresses nuclear translocation of SREBP1, and inhibits the transcription of its direct target gene BCAT2. We further revealed BCAT2 as a suppressor of ferroptosis by regulating intracellular glutamate level (Figure5). Importantly, the combination of sulfasalazine and sorafenib has synergistic effect in inhibiting BCAT2 expression as well promoting ferroptotic cancer cell death *in vitro* and in a couple of animal models including in subcutaneous pancreatic cancer model, orthotopic liver cancer model, as well as PDX hepatic carcinoma model. Of importance, BCAT2 has also shown the potential as a sensitive biomarker to evaluate drug responses in these preclinical cancer models.

The finding that BCAT2 controls the ferroptosis is in accordance with the concept that amino acids play a crucial role in ferroptosis(Gao & Jiang, 2018). BCATs are key metabolic proteins catalyzing the reversible transamination of BCAAs to their respective a-ketoacids (BCKAs) and glutamate, responsible for the production of 30% of *de novo* brain glutamate(LaNoue, Berkich et al., 2001, Lieth, LaNoue et al., 2001). The metabolism of glutamate is tightly linked to the regulation of ferroptosis(Angeli et al., 2017, Gao et al., 2015). It is important to note that system Xc^−^ function is regulated by glutamate levels since glutamate is exchanged for cystine in a 1:1 ratio by system Xc^−^. Accordingly, high extracellular concentrations of glutamate block system Xc^−^ activity, inhibit cystine uptake, and drive ferroptosis(Dixon et al., 2012). In contrast, high level of intracellular glutamate in our *in vitro* experiments derived from BCAT2 driven *de novo* synthesis of glutamate, which may consequently enhance system Xc^−^ activity, boost cystine uptake, and inhibit ferroptosis. This protective effect of BCAT2-intracellular glutamate metabolism is consistent with the fact that there are decreased extracellular brain glutamate levels protected system Xc^−^ knockout mice from neurotoxic injury(Massie, Schallier et al., 2011).

BCAAs are nitrogen donors for the synthesis of not only glutamate but also glutamine, but the role of glutamine in ferroptosis is complex. Glutamine is degraded through its specific metabolism, glutaminolysis. When glutaminolysis is inhibited or glutamine is depleted, cystine starvation and blockage of cystine import fail to induce ROS accumulation, lipid peroxidation, and ferroptosis, indicating that glutaminolysis fuels ferroptosis. In line with this observation, a-ketoglutarate (a-KG), another product of glutaminolysis besides glutamine, can replace glutamine to induce ferroptosis (Gao et al., 2015). We speculate that BCATs catalyze BCAAs-BCKAs shuttle for the synthesis of glutamate, leading to a reduction of intracellular level of a-KG, which may be another reason to induce ferroptosis. Knockdown BCAT2 has no effect on the expression of SLC7A11.Therefore, the exact relationship between BCAT2 and system Xc^−^ need deeper investigation in the future work.

Liver cancer is the third leading cause of cancer deaths worldwide, and standard chemotherapy has not been effective in most patients with liver cancer, doctors have been looking at targeted therapies. Sorafenib is the only multikinase inhibitor as the first-line treatment proven to prolong overall survival of unresectable hepatocellular carcinoma (Tovoli, Ielasi et al., 2019). However, the overall survival in patients from the Asia-Pacific region taking sorafenib was just 6.5 months with low response rate of 2%. Lenvatinib is recently approved as an alternate multikinase agent for advanced hepatocellular carcinoma if sorafenib stops working, but its overall survival superiority over sorafenib was not achieved in a recent phase 3 clinical study. In this study we found sulfasalazine alone, or in combination with sorafenib, function in ferroptosis-inducing therapies. These findings are consequential since sulfasalazine is an anti-inflammatory drug which has already been used extensively in chronic, long term therapy of inflammatory bowel disease, guaranteeing its safety profiles both in adults and children(Scaioli, Sartini et al., 2017). Based on our results, sulfasalazine might be a potential new treatment option for advanced liver cancer, as well as other unresectable cancer types. Due to the expression changes in the treatment, BCAT2 may be one of the most sensitivity targets and its expression can be useful as a marker predicting response to sorafenib and sulfasalazine combination treatment. However, this hypothesis should be evaluated in patient data.

Taken together, our data demonstrate that inhibiting intracellular glutamate synthesis could serve as a good strategy for inducing ferroptosis in cancer contexts. This is supported by our finding that sulfasalazine collaborates with sorafenib to downregulate BCAT2 and consequently intracellular glutamate. Our work also suggests a mechanism for cell lethality involving the regulation of *de novo* synthesis of glutamate as crucial process in liver cancer cells. We suggest that the protein or mRNA level of BCAT2 may be used to predict the responsiveness of cancer cells to future ferroptosis-inducing therapies. We also propose that highly specific BCAT2 inhibitors could provide an effective therapy for a meaningful fraction of cancer patients.

## Materials and Methods

### Cell Culture and Reagents

Aspc-1, HepG2, Panc02 and H22 cell cell lines were obtained from the KeyGEN Biotechnology Company (China). HT1080 and SW480 were obtained from the FuHeng BioLogy Company (China). HT1080 cancer cells were cultured in Eagle’s Minimum Essential Medium (EMEM) supplemented with 10% fetal bovine serum (FBS), glutamine (2 mM), penicillin (100 U/ml) and streptomycin (0.1 mg/ml). SW480, Aspc-1, HepG2, Panc02 and H22 were cultured in high Dulbecco’s Modified Eagle’s Medium (DMEM) supplemented with 10% FBS, L-glutamine (4 mM), and penicillin(100 U/ml) and streptomycin (0.1 mg/ml). All cell lines were maintained in a humidified atmosphere containing 5% CO_2_ at 37℃ and tested for mycoplasma prior to the commencement of experiments. Unless otherwise indicated, cell culture medium was changed every three days, and cells were passaged using 0.05% trypsin/EDTA. Erastin(HY-15763), sorafenib(HY-10201), sulfasalazine(HY-14655), ferrostatin-1(HY-100579), Z-VADFMK(HY-16658), AICAR(HY-13417), BafA1(#HY-100558), and Necrosulfonamide (HY-100573) were purchased from MedChemExpress (MCE, USA). Compound C (#ab120843) were purchased from Abcam. Deferoxamine mesylate (#D9533) were purchased from Sigma Aldrich.

### CRISPR activation (CRISPR a) screen

HepG2 cells were infected with lentivirus encoding Cas9-VPR enzyme and selected with 2μg/ml puromycin. Briefly, 4.343 × 10^7^ cells were infected with human CRISPRa sgRNA lentivirus library targeting 2320 genes with about 13030 sgRNA at a low MOI (0.3). After 48 h, the infected cells were selected with 800 μg/ml G418 for 72 h. Cells were equally split into 2 samples (at least 1.3×10^7^/sample). One sample was treated with 10 μM Erastin for 16 h and changed back to DMEM once the 3 days for 3 rounds compared with another untreated sample as control. Genomic DNA was extracted and sgRNA were amplified by PCR. The resulting PCR products were sequenced by Illumina Hiseq 4000 and evaluated based on the known sgRNA targets sequence.

### Cell viability assay

Tumor cells were collected and seeded into 96-well plates. After adhesion, cells were treated with the different ferroptosis inducers or inhibitors. To determine the effect of treatment on cell viability, Cell Counting Kit-8(CCK8, #CA1210, Solarbio) was carried out according to the manufacturer’s instructions. Absorbance at wavelengths of 450 nm was measured. The percentage difference in reduction between treated and control cells was calculated. After calculation, the viability of control cells was 100% and all others were normalized to control and shown as relative cell viability (%).

### Western blot assay

Protein was quantified using the bicinchoninic acid (BCA) assay (Thermo Fisher Scientific, #23225). Western blotting assay was performed as described previously(Chen, Cheng et al., 2019). Antibodies were as follows: anti-human BCAT2(CST, #9432S, 1:1000), anti-human BCAT1(Abcam, #ab197941, 1:1000), anti-human Phospho-AMPKα (Thr172)(CST, #2535, 1:1000), anti-human AMPK(CST, #5831, 1:1000), anti-human SREBP1(Santa Cruz, #SC-13551, 1:200), anti-human NCOA4(Abcam, #ab86707, 1:1000), anti-human ATG7(CST, #8558, 1:1000), anti-human SLC7A11(Abclonal, #A13685, 1:1000), anti-human GPX4(Abcam, #ab41787, 1:1000), and β-tubulin (Abcam, #ab6046, 1:1000). Secondary antibody (either anti-rabbit or anti-mouse) was purchased from Boster Biotechnology Company (China). The blots were analyzed using the software ImageJ (Version 1.80, NIH, USA)

### Quatitive real time polymerase chain reaction assay (qRT-PCR)

Total RNA was extracted using TRIzol (Invitrogen) according to the manufacturer’s instructions. RevertAid First-Strand cDNA Synthesis Kit (Thermo, Waltham, MA, USA) was performed for reverse transcription according to the manufacturer’s specification. Subsequently, SYBR Green-based real-time PCR was performed in triplicate using SYBR Green master mix (Thermo Fisher Scientific) on an Applied Biosystems StepOnePlus real-time PCR machine (Thermo Fisher Scientifc). For analysis, the threshold cycle (Ct) values for each gene were normalized to expression levels of GAPDH. Analysis was performed using the Bio-Rad CFX Manager software. The primers, which were synthesized and desalted from Sigma-Aldrich, are shown in Table1.

### Chromatin immunoprecipitation(ChIP)

ChIP was performed according to the protocol of the chromatin immunoprecipitation assay kit. Briefly, cells were pretreated erastin, sorafenib, and sulfasalazine, respectively, and then cross-linked in 3.7% formaldehyde for 15min, quenched with glycine for 5min, and lysed with SDS lysis buffer. Chromatin was sheared by sonication, and lysates were precleared with salmon sperm DNA/protein A agarose slurry for 1h and incubated with rabbit IgG (Santa Cruz) or SREBP1 antibody in the presence of protein A agarose beads overnight. After sequential washes of the agarose beads and eluted, the elutes were heated at 65°C for 4 h to reverse the cross-linking and treated with RNase A for 30 min at 37°C, followed by treatment with proteinase K for 1 h at 45°C to remove RNA and protein. DNA was recovered, eluted, and then assayed using PCR. The ChIP primers were purchased from Qiagen (EpiTect ChIP PCR assay) and used for qPCR analysis: BCAT2.

### RNAi and gene transfection

Cancer cells were seeded in 6-well plates at a density of 1 × 10^5^ cells/well to achieve a confluence of 70-80% overnight. To generate BCAT2 knockdown cells, cells were transfected with 10 nM of shRNA against BCAT2 and negative control shRNA (Suzhou Ribo Life Science CO., Ltd, China), respectively. Transfection was performed with Lipofectamine 2000 (Invitrogen) according to the manufacturer’s instructions. The specific shRNA sequences are listed in Table 2.

**Table 1.**
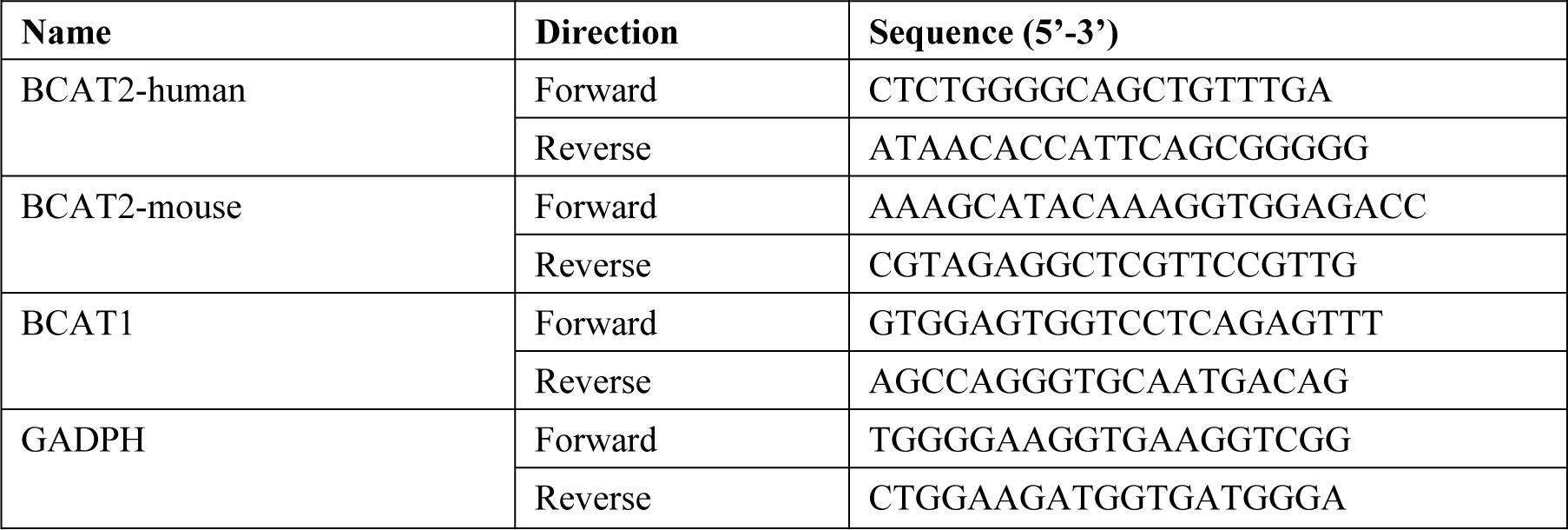
Sequences of primers used for qRT-PCR.

**Table 2.**
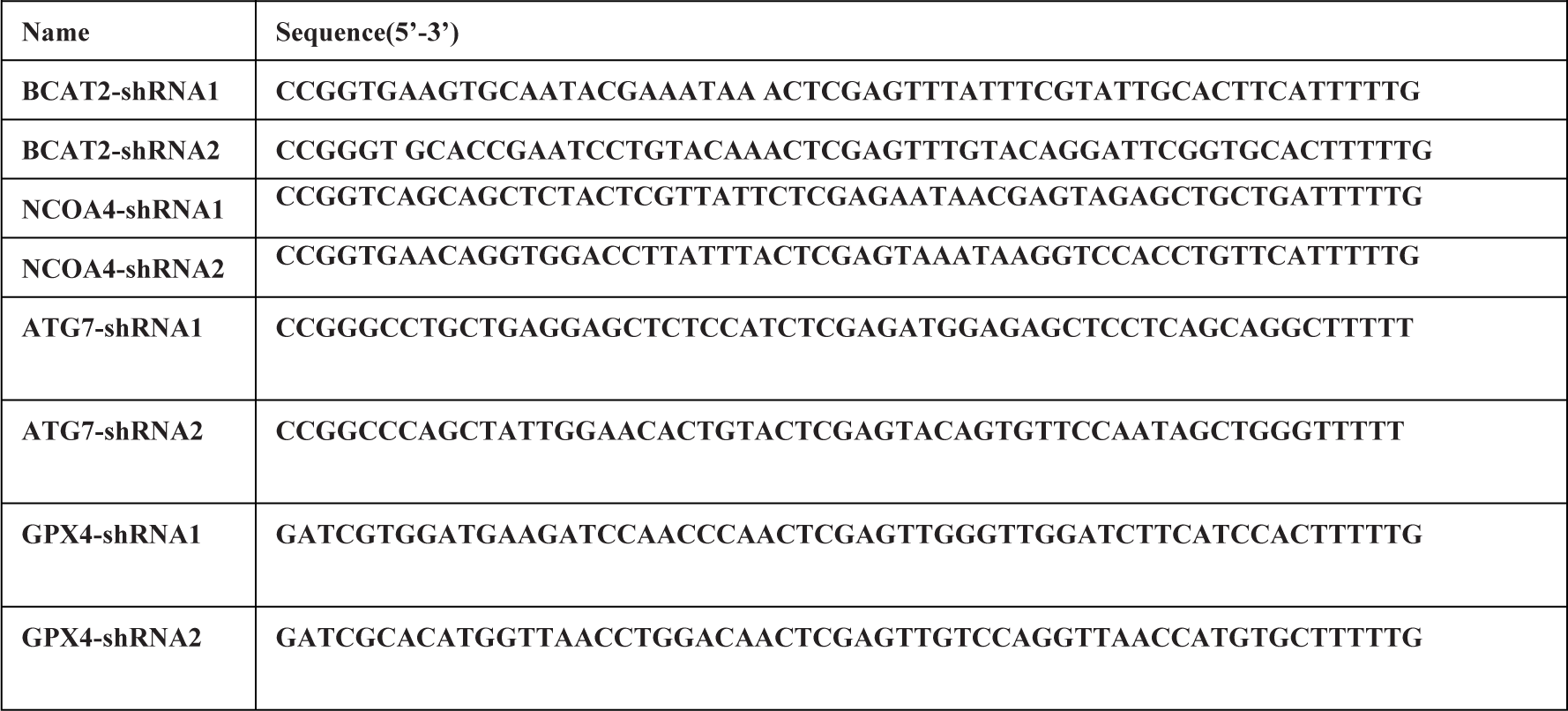
Sequences of shRNAs.

For establishing the stable sh-BCAT2 cancer cells, the lentiviral packaging kit was purchased from Open GeneCopoeia. Lentivirus carrying BCAT2-shRNA was packaged in 293T cells and concentrated from the supernatant, as instructed by the manufacturer’s manual. Stable cell lines were established by infecting lentivirus into cancer cells followed by puromycin (1μg/ml) selection for 10-14 days. These established stable cell lines were maintained in DMEM containing 10% FBS and puromycin (0.75μg/ml) for further experiments.

### BCAT2 overexpression experiment

The mammalian expression plasmid pLVx-BCAT2-Flag (FC-324) was purchased from the Fubio Company (Suzhou, China). Cells were transfected with the stated constructs according the manufacturer’s instructions (Invitrogen, China). Cancer cells were seeded in a 96 well dish at a density of 2000 cells/well. The following day, cells were infected with the vector described above. Cells were infected at an M.O.I. of ∼1 in media containing 8 µg/mL polybrene and spun at 1000 rpm for 1 h at room temperature. The next day, virus-containing medium was removed and replaced with medium containing 10 µg/mL puromycin. After 24 h, cells were treated with various agents for further use.

### Iron assay

Intracellular ferrous iron level was determined using the iron assay kit (Abcam, ab83366) according to the manufacturer’s instructions.

### Lipid peroxidation assay

The relative MDA concentration in cell or tumor lysates was assessed using a Lipid Peroxidation (MDA) Assay Kit (Abcam, #ab118970) according to the manufacturer’s instructions. Briefly, MDA in the sample reacts with thiobarbituric acid (TBA) to generate a MDA-TBA adduct. The MDA-TBA adduct can be quantified colorimetrically (OD=532 nm). C11-BODIPY dye (Thermo Fisher Scientific) was used to detect lipid peroxidation in cells. Oxidation of the polyunsaturated butadienyl portion of the dye results in a shift of the fluorescence emission peak from ∼590 nm to ∼510 nm.

### Glutathione (GSH) assay

The relative GSH concentration in cells was assessed using a GSH/GSSG Ratio Detection Assay Kit (Abcam, #ab205811) according to the manufacturer’s instructions. Briefly, whole cell were diluted to 1: 80 for GSH analysis, serial dilution of GSH and GSSG stock standards were prepared as standards. A one-step fluorimetric reaction of samples with respective assay buffer and probes were incubated for 30 min. The yellow product (5-thio-2-nitrobenzoic acid) was measured spectrophotometrically at 412 nm

### Glutamate release assay

The relative release of glutamate from cells into the extracellular medium was detected using an Amplex Red glutamate release assay kit (Thermo Fisher Scientific). Glutamate release was first normalized to the total cell number determined with the CCK8 kit at the end of the experiment, and then values were expressed as a percentage of no-treatment controls.

### Intracellular glutamate assay

The relative intracellular glutamate was detected using a Glutamate Assay Kit (Abcam, #ab83389) according to the manufacturer’s instructions. The intracellular glutamate first normalized to the total cell number determined with the CCK8 kit at the end of the experiment, and then values were expressed as a percentage of no-treatment controls.

### TEM imaging

To observe the subcellular structure, cancer cells treated with various agents were harvested and fixed with 2.5% glutaraldehyde in 0.1 M sodium cacodylate buffer for 24 h, and subsequently fixed in 1% osmium tetroxide for 2 h. Specimens were dehydrated in a graded series of acetone and embedded in epoxy resin. After ultramicrotomy, ultrathin sections were stained with uranyl acetate for 15 min and modified with lead citrate for 5 min. Finally, the subcellular structure was observed by TEM using a JEOL JEM-2100 microscope (JEOL Ltd. Japan).

### Colony forming assay

Cell growth of shRNA-treated cell lines was assayed through crystal violet staining. For colony formation assays, 2000 cells were seeded in 6-well plates for 14 days. Cells were fixed with 80% methanol and stained with crystal violet solution overnight. All experiments were performed in triplicate.

### Immunofluorescence

HepG2 cells stably expressing GFP-LC3 were grown on glass cover slips in a six-well plate. After 24 h, cells were treated with various agents for 24 h. The cells were then washed with PBS and fixed in 3.7% paraformaldehyde for 10 min at 37°C. The number of GFP-LC3 dots was then detected using a confocal fluorescence microscope.

### Immunohistochemistry analysis

Paraffin-embedded tissues were sectioned for immunohistochemical (IHC) analysis. For IHC, samples were fixed in 10% formalin and embedded in paraffin wax. Next, 3 mm sections were cut from the paraffin blocks for IHC analysis. The sections were stained with mouse rabbit-BCAT2 (Abcam, #ab95976, 1:200) at 4°C overnight. All the sections were cover slipped with neutral balsam and viewed under an Olympus microscope and analyzed using Image J software. The final result for each case was the average score of all visual fields.

### Xenograft tumor models

Animal studies were approved by the Committee on the Use of Live Animals for Teaching and Research of the Jiangsu University. Female C57BL/6 mice (purchased from The Compare Medicine Center, Yangzhou University, China), age 4 weeks, were health checked daily throughout the experiment and kept on a regular 12 h light and dark cycle with normal diet in a pathogen-free barrier facility. 1 × 10^6^ BCAT2 overexpression and control Panc02 cancer cells were implanted subcutaneously into the right dorsal flanks of C57BL/6 mice (five mice per group), respectively. When the tumors reached a volume of 50-100 mm^3^, the mice were treated with or without erastin (40 mg/kg) every two days for 2 weeks. Due to the low solubility and poor metabolic stability, erastin was administration by the intratumoral injection. The tumor volume and growth speed was monitored every two days until the end point at day 14.

To investigate the role of combination sorafenib with sulfasalazine inducing ferroptosis, 1 × 10^6^ Panc02 were implanted subcutaneously into the right dorsal flanks of C57BL/6 mice. When the tumors reached a volume of 50-100 mm^3^, mice were randomly divided into four groups (five mice per group) and treated with DMSO (control), sorafenib(10 mg/kg), sulfasalazine (100 mg/kg), or the combination of these drugs at the indicated doses by intraperitoneal injection every two days for two weeks.

The tumor volume and growth speed was monitored every two days until the end point at day 14.

To generate orthotopic tumors, forty C57BL/6 mice were surgically implanted with 1 × 10^6^ H22 cells into left lobe of livers. One week after implantation, mice were randomly allocated into four groups (ten mice per group) and treated with the following agents: (i)DMSO; (ii)sorafenib (30 mg/kg); (iii) sulfasalazine (300 mg/kg); or (iii) sorafenib(30 mg/kg) + sulfasalazine (300 mg/kg) by intragastrical(i.g.) administration every two days for three weeks. Animal survival was calculated every day for 2 months. Fresh tumor tissue weight was immediately accessed following harvest.

### PDXs and In Vivo Experiments

NSG (NOD. Cg-*Prkdc^scid^ Il2^rgtm1Wjl^*/SzJ) mice were purchased from the BEIJING IDMO Co., Ltd. and maintained in Animal Center of Jiangsu University in compliance with the Guide for the Care and Use of Laboratory Animals (NIH Publication No. 85–23, revised 1996). The experimental protocols were approved by the Committee for Ethical Affairs of Jiangsu University (Zhenjiang, China), and the methods were carried out in accordance with the approved guidelines.

Serial passaging of the PDX was carried out by implanting small fragments of the liver tumor subcutaneously into dorsal flanks of NSG mice. Experiments were performed using PDX tumors passages 4 and 5. PDXs mice were randomly allocated into four groups (five mice per group) and treated with the following agents: (i)DMSO; (ii)sorafenib (10 mg/kg); (iii) sulfasalazine (100 mg/kg); or (iii) sorafenib(10 mg/kg) + sulfasalazine (100 mg/kg) by intraperitoneal every two days for 28 day. The body weight, tumor volume and growth speed was monitored every two days until the end point at day 40. Tumor weight was fresh tumor tissues from all the mice. Tumor tissues were stored for MDA assay, qRT-PCR, west blotting and immunohistochemistry analysis.

### Patient selection

The Cancer Genome Atlas (TCGA) database: https://tcga.xenahubs.net/download/TCGA.LIHC.sampleMap/HiSeqV2.gz.

Hepatocellular carcinoma gene expression by RNAseq (IlluminaHiSeq percentile) including 390 hepatic carcinoma patient specimens was utilized to further analyze the relationship between BCAT2, GPX4, NCOA4, TP53, BECN1, NRF2, and SLC7A11. High and low groups were defined as above and below the mean respectively.

### Statistics

All data are presented as the mean ± standard error of the mean (SEM). Linear regression and F testing were used to determine correlation between BCAT2, GPX4, NCOA4, TP53, BECN1, NRF2, and SLC7A11 in hepatocellular carcinoma. The significances of differences between groups were analyzed using Student’s t tests, one-way analysis of variance (ANOVA) or two-way ANOVA. *P* < 0.05 was considered to reflect a statistically significant difference. All the experiments were repeated at least three times.

## Author contributions

Haitao Zhu, Fang Cheng, and Dongqing Wang served as corresponding authors and organized the study. Kang Wang, Zhengyang Zhang, and Tsai Hsiang-i performed the experiments. Yanfang Liu, Ming Wang, Hongbo Chen and Lian Song analyzed the data. Xiongfeng Cao and Zhanxue Xu organized the figures. Haitao Zhu and Aihua Gong drafted the manuscript. All authors read and approved of the final manuscript.

## Acknowledgements

This study was supported by grants from the National Natural Science Foundation of China (grant numbers 81502663, 81702750), the Social Development Foundation of Jiangsu Province (grant number BE2018691, BK20191223), Young Medical Talents of Jiangsu (grant number QNRC2016833), Six talent peals project of Jiangsu Province (grant number WSW-039), Six for one project of Jiangsu Province (grant number LGY2018093), and the Social Development Foundation of Zhenjiang City (grant numbers SH2018063, SH2018031), the Basic Research Project of Shenzhen (grant numbers JCYJ20170818164756460, JCY J20180307154700308).

